# Alphavirus nsP2 protease structure and cleavage prediction: Possible relevance to the pathogenesis of viral arthritis

**DOI:** 10.1101/2022.01.22.477317

**Authors:** Lucas Prescott

## Abstract

Alphaviruses are a diverse genus of arboviruses capable of infecting many vertebrates including humans. Human infection is common in equatorial and subtropical regions and is often accompanied by arthralgia or encephalitis depending on viral lineage. No antivirals or vaccines have been approved, and many alphavirus lineages have only recently been discovered and classified. Alphavirus nsP2 protease is an important virulence factor yet is commonly thought to be a simple papain-like protease which only cleaves viral polyproteins. Here, I reveal novel molecular mechanisms of these proteases via sequence and predicted structure alignment and propose novel cellular mechanisms for the pathogenesis of viral arthritis by predicting which human proteins are likely cleaved by these proteases. In addition to the known primary cysteine mechanism in all alphaviruses and a secondary serine mechanism documented in chikungunya virus (CHIKV), I discovered secondary cysteine and threonine mechanisms exist in many other alphaviruses and that these secondary mechanisms coevolve with their viral polyprotein cleavages. As for cleavage prediction, neural networks trained on 93 different putative viral polyprotein cleavages achieved a Matthews correlation coefficient of 0.965, and, when applied to the human proteome, predicted that hundreds of proteins may be vulnerable. Notable pathways likely affected by cleavages include the cytoskeleton and extracellular matrix, antiproteases, protein translation/folding/glycosylation/ubiquitination, cellular differentiation, inflammation, and vesicle trafficking, hinting that this viral protease is a more important virulence factor than previously believed.

## Introduction

Alphavirus genomes contain two open reading frames encoding non-structural and structural polyproteins. Although the structural polyprotein is proteolytically processed by the capsid protein and host furin and signal peptidases, the non-structural polyprotein is processed typically by a cysteine protease contained within nsP2 (nsP2pro). A cleft for substrate binding exists between nsP2’s C-terminal protease and S-adenosyl-L-methionine-dependent methyltransferase (MTase)-like subdomains connected by a flexible linker, and long-range interactions with nsP2’s N-terminal helicase[1, 2] or any nsP3 domains before its separation are important for polyprotein processing and virulence yet remain poorly characterized.[3, 4] CHIKV nsP2pro has been found to not only contain a papain-like cysteine mechanism, but also an adjacent serine with similar activity.[5] This mechanism has not yet been found in any other alphaviruses, but it likely dramatically affects the stability, activity, and selectivity of nsP2pro. Additionally, a single mutation (N475A) near the N-terminus of the protease subdomain was found to cause the flexible N-terminal residues to occupy the cleft and inhibit catalysis.[6] This mutation does not, however, exist in any natural variants, and this study was not performed on full-length nsP2. The nuclear localization signal and RNA-binding helicase determining nuclear and virion localization of nsP2 likely also drive nsP2pro’s selective pressures and multiple activities.[7] In addition to these subdomain interactions, most alphaviruses contain a stop codon at the end of nsP3 which is read through in 5-20% of polyproteins.[7, 9] Depending on this termination suppression, nsP2pro cleaves either two or three sites within the non-structural polyprotein with kinetic rates varying up to 25 fold[10, 11] and, as with many viral proteases, is expected to cleave many host factors. To my knowledge, the antiviral TRIM14 is the only host protein experimentally verified to be cleaved by an alphavirus nsP2pro (Venezuelan equine encephalitis virus (VEEV) and somewhat by other New World alphaviruses).[12]

Due to the few cleavages per polyprotein and the continually expanding taxonomy of alphaviruses, few viral proteases or their cleavages have been characterized.[13] Following the successful application of machine learning methods to other viral proteases for cleavage prediction in human proteins,[14–16] computational analysis of alphavirus proteases will likely be an important step toward discovering additional therapeutic targets.

## Methods

### Data Set Preparation

A complete, manually reviewed human proteome containing 20,350 sequences (not including alternative isoforms) was retrieved from UniProt/Swiss-Prot (proteome:up000005640 AND reviewed:yes).[17] All polyprotein sequences within the family *Togaviridae* were collected from GenBank,[18] and 93 different cleavages were manually discovered using the Clustal Omega multiple sequence alignment server.[19–21] Similar cleavages are discoverable in divergent species within the order *Martellivirales* but were not included here because none infect animals. The next closest species that can infect humans are rubella (RUBV) and hepatitis E (HEV) viruses within the broader class *Alsuviricetes,* but their non-structural proteases have drastically different structures and activities than those within *Togaviridae* and were therefore also omitted. All unbalanced positive cleavages were used for subsequent classifier training in addition to all other 5,461 uncleaved alphavirus sequence windows with glycines in the P2 position, totaling 5,554 samples.

### Protease Structure Prediction and Analysis

AlphaFold was used to predict the structures of alphavirus nsP2 sequences (only the C-terminal protease and MTase-like subdomains).[22] Predicted backbones were nearly identical to experimental data, and predicted catalytic dyad side chain distances ranged from 4 to 8 Å. Although AlphaFold does not have the ability to accurately predict the impact of single missense mutations on protein structures,[23, 24] this set of predicted structures and simulations, albeit on often nearly identical sequences, serves as a starting point to understanding the diversity of mechanisms within alphaviruses. CABS-flex was used to predict alternate conformations and flexibilities,[25] and zinc binding prediction was performed with ZincBind.[26] Molecular graphics and analyses were performed using UCSF ChimeraX, developed by the Resource for Biocomputing, Visualization, and Informatics at the University of California, San Francisco, with support from National Institutes of Health R01-GM129325 and the Office of Cyber Infrastructure and Computational Biology, National Institute of Allergy and Infectious Diseases.[27]

### Cleavage Prediction and Analysis

As in my previous work on 3CLpro and PLpro,[14, 15] sequence logo-based logistic regression and naïve Bayes classification and physiochemical and one-hot encoded neural networks (NNs) were used for cleavage prediction.[28] To reduce any potential false positives, only proteins expressed in relevant cell types with cleavages with agreement between all five NN replicates and with total solvent-accessible surface areas (SASAs) of more than 100 Å^2^ between positions P5 and P5’ were reported. SASAs were calculated from AlphaFold predicted human protein structures[22] with the FreeSASA package.[29] Multiple synovial fluid and associated cell type proteomes and transcriptomes were compiled and cross-referenced to remove cleavages irrelevant to arthritic pathogenesis.[30–35] Predictions of the effects of cleavage on subcellular localizations were performed using the DeepLoc server.[36] All training data, prediction methods, and results can be found on GitHub (https://github.com/Luke8472NN/NetProtease).

## Results

Although alphavirus proteases are diverse and not necessarily only papain-like, their cleavages resemble those of coronavirus papain-like protease (PLpro) but not of papain itself (Figure 1).[37] The repeated glycines and alanines in alphavirus cleavage positions P2, P1, and P1’ were, however, easier to align than coronavirus PLpro cleavages. Dimensionality reduction of putative cleavages clustered by order within the polyprotein much more than by lineage (except for previously discovered cleavages in tymoviruses[38]), indicating that all cleavages within *Togaviridae* but not *Alsuviricetes* can be combined into a single data set to train machine learning models to apply to human sequences (Figure 2). Although no cleavages from species outside *Togaviridae* were included for training here, it is noteworthy that turnip yellow mosaic virus (TYMV) protease is but HEV and RUBV proteases are not structurally related to alphavirus nsP2pro. TYMV protease includes an equivalent catalytic cysteine helix and activity-tuning histidine flexible loop[39] yet does not contain an MTase-like domain to form a cleft as in alphaviruses. In addition to this more accessible active site, TYMV protease includes two hydrophobic patches required for interaction with ubiquitin for its deubiquitinating activity (Figure 3).[40]

**Figure 1:**
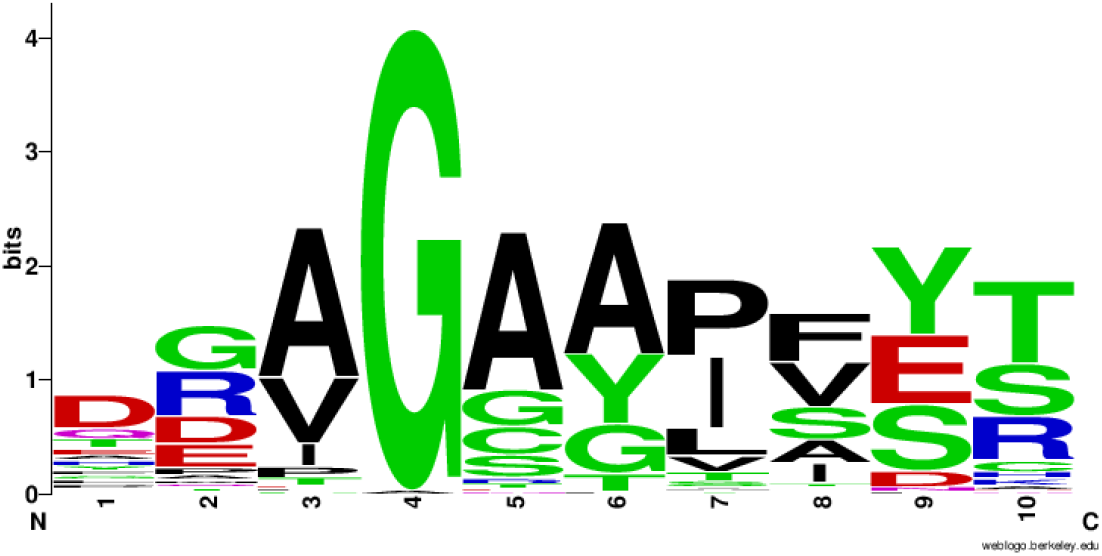
Sequence logo for all 93 putative cleavage sites. [41]

**Figure 2:**
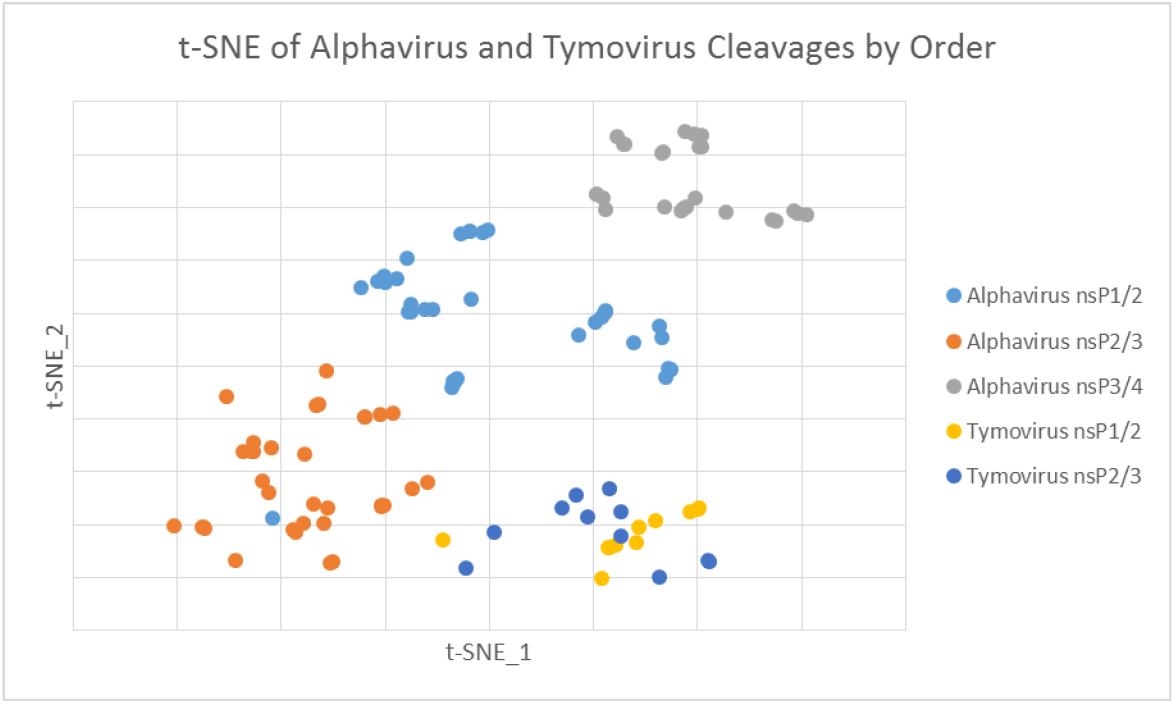
One-hot encoded cleavage t-SNE colored by order within the polyprotein and by genus. [42]

**Figure 3:**
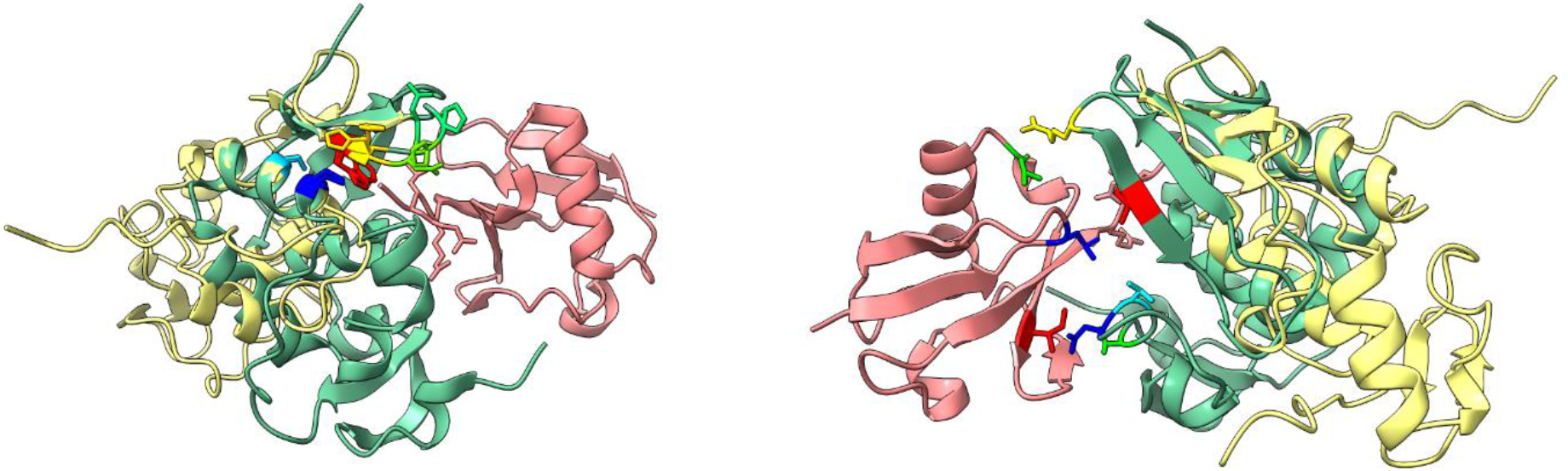
(A) Structural similarity between CHIKV and TYMV proteases near their active sites but (B) not in ubiquitin binding regions. Only TYMV protease binds ubiquitin’s I36 and I44 hydrophobic patches and its L8 loop. Tan ribbon is CHIKV, green ribbon is TYMV, red ribbon is ubiquitin (PDB code 6YPT).[27, 40]

Alignment of all known alphavirus proteases (Figure 4) indicated that, in addition to a primary cysteine mechanism in all alphaviruses and a secondary serine mechanism found in at least CHIKV,[5] some proteases have secondary cysteine or threonine mechanisms. These secondary mechanisms may restrict or extend the possible acidic residues aligning and polarizing the catalytic histidine,[43] and serine and threonine mechanisms may extend catalytic activity to higher pH.[44] By investigating how substrate sequences, particularly the P1 residue, coevolve with these different mechanisms, multiple functional hypotheses can be proposed: (1) cleavage after a P1 cysteine is most efficient when the secondary catalytic residue is another cysteine or serine, possible for an inert secondary alanine, and least efficient for a secondary threonine, (2) a secondary threonine is required for cleavage after a P1 serine, and (3) an inert secondary alanine is required for cleavage after a P1 arginine.

**Figure 4:**
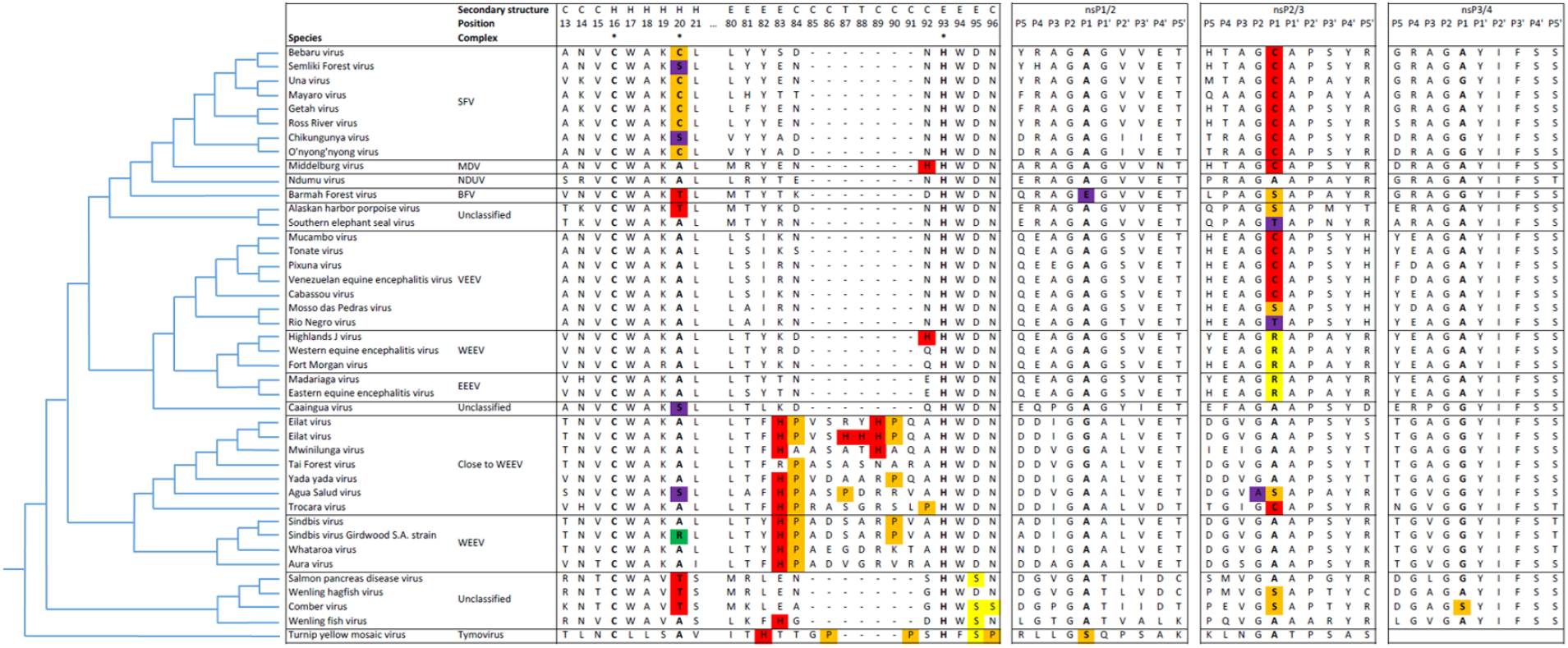
(A) Cladogram and multiple sequence alignment of the catalytic dyad and flexible loop of alphavirus and TYMV proteases and (B) their respective aligned cleavages.

Near the catalytic histidine, other histidines or related positively charged amino acids in the longer flexible loops of Western equine encephalitis complex and related viruses are close enough in proximity with each other that they may bind ordered water molecules as in other proteases[45] or metal ions in multiple conformations.[26] Unlike RUBV cysteine protease[46] and hepatitis C virus (HCV) NS3 serine protease[47] which require metal ions in either a structural or catalytic (as in metalloprotease) role for activity, metal ions are known to inhibit CHIKV protease.[48] In these alphavirus proteases, metal binding may disrupt normal histidine alignment with aspartic acid[49] and aim its protonated side toward the catalytic cysteine, serine, or threonine, preventing the charge relay mechanism required for proteolysis (Figure 5A). Additionally, metal binding to another histidine in the center of the loop may aid its flexing backward to allow substrate loading. In some divergent, unclassified alphavirus proteases, the adjacent aspartic acid is replaced with serine, but in these cases there is always another nearby potential metal-binding residue (glutamic acid, aspartic acid, or another histidine)(Figure 5B).

**Figure 5:**
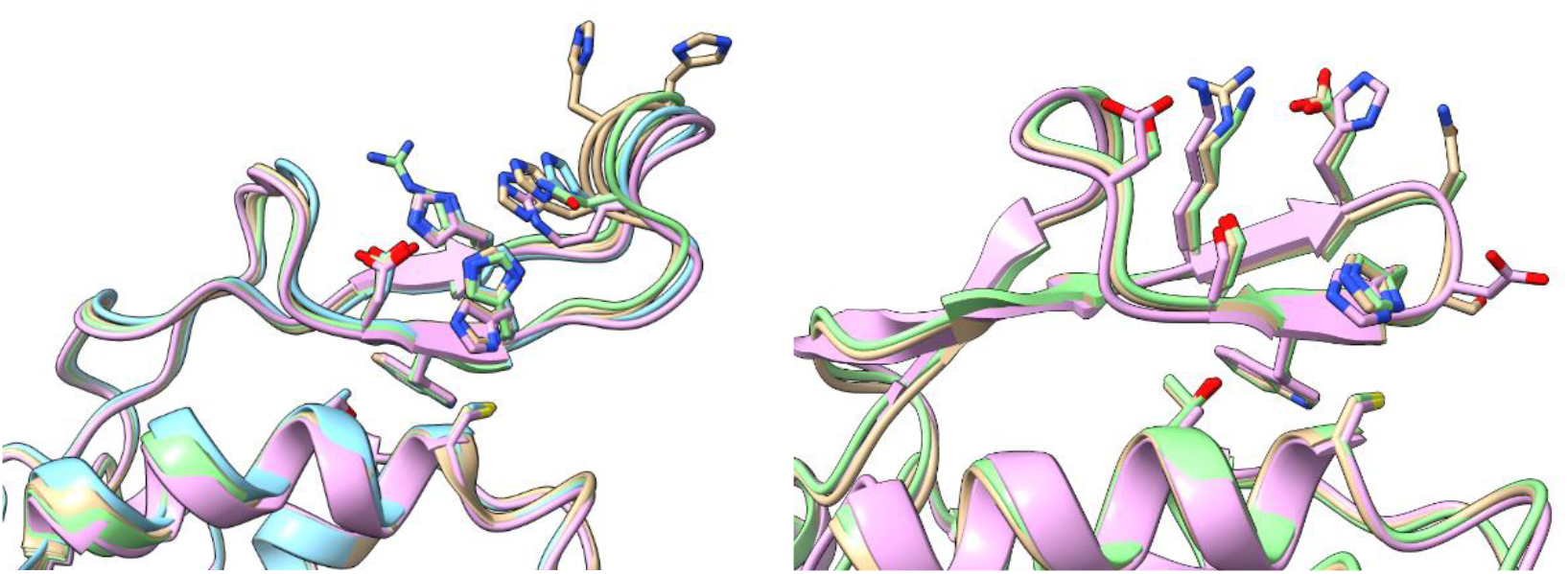
(A) Proposed metal binding histidines and adjacent aspartic acid guiding charge relaying in proteolysis. Tan ribbons are Eilat viruses, one of which includes five histidine residues,[50] purple ribbon is Agua Salud virus (ASALV), green ribbon is Tai Forest virus, and blue ribbon is Mwinilunga virus. (B) Proposed alternative metal binding residues in divergent proteases. Tan ribbon is Salmonid virus, purple ribbon is Wenling fish virus, and green ribbon is Comber virus).[27]

To my knowledge, no P2 glycine substitutions have been discovered in alphavirus cleavages, so it is noteworthy that ASALV[51] nsP2/3 cleavage (DGVAS^APAYR in MK959114.1 and MK959115.1) contains an alanine in this position (Figure 4). No sequence features obviously correlated with this substitution, and ASALV protease’s predicted structure is extremely similar to those of related alphavirus proteases, indicating that other alphavirus proteases may also cleave alanine-containing substrates albeit possibly with suboptimal kinetics. Tryptophan is typically thought not to be directly involved in the active site, yet it appeared here to obstruct the secondary catalytic mechanism in some conformations (Figure 6A). In addition to the flexibility of the catalytic dyad, the size and flexibility of the variable loop between the protease domain β1 and β2 strands and its interaction with the MTase-like domain loop between β7 strand and α9 helix (Figures 6B and 6C) likely determine the rate of substrate loading into the cleft and therefore cleavage kinetics.[52] Deletion of the exposed and most proximal MTase-like domain residue (typically leucine, phenylalanine, or tryptophan) and sharper backbone twisting by subsequent prolines in divergent alphaviruses may also widen the gap between these two loops and affect substrate loading or may allow serine to better fit in the P1 pocket instead of the more common alanine (Figure 6D). Sodium[53, 54] or other salt binding within this cleft may affect structure and substrate binding, and these deep cleft residues are not conserved between alphaviruses. No matter the width of this cleft, the nsP2/3 site remained over 40 Å away from the active site, supporting this site’s proposed *trans* cleavage.[3] Even Salmonid virus’ large insertions at the C-terminus of its nsP2 MTase-like domain and between its nsP3 macro and zinc-binding domains did not affect this distance.

**Figure 6:**
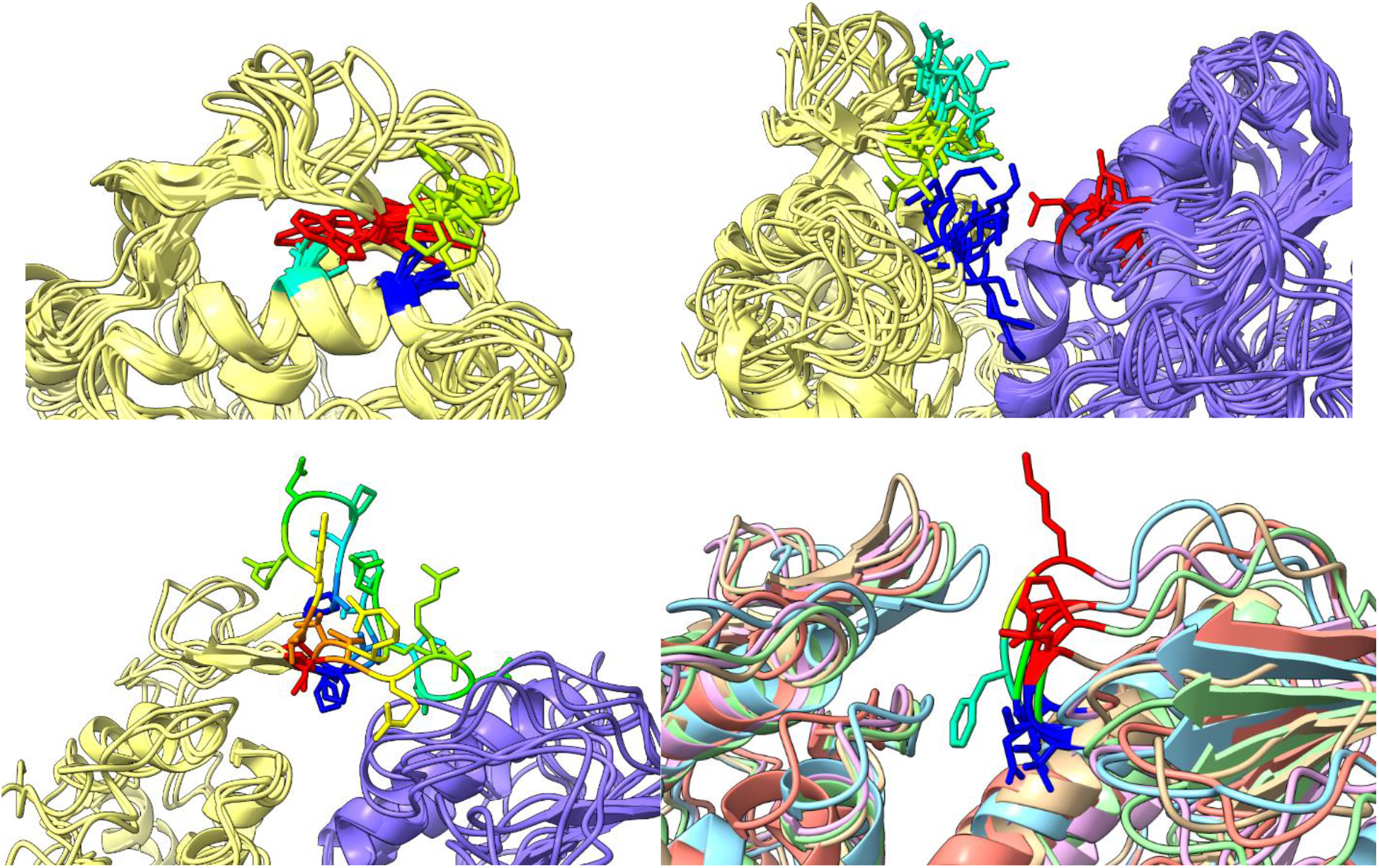
(A) CABS-flex ensemble of CHIKV nsP2pro flexible catalytic core and (B) flexible loop between protease subdomain (tan ribbon) β1 and β2 strands interacting with loop between MTase-like domain (purple ribbon) β7 strand and α9 helix. (C) ASALV insertion within flexible loop.[25] (D) Similarity between interacting MTase-like domain loops with noteworthy deletions. Tan ribbon is Alaskan harbor porpoise virus, blue ribbon is Salmonid virus, purple ribbon is Wenling fish virus, green ribbon is Wenling hagfish virus, and red ribbon is Comber virus.[27]

As with both coronavirus protease cleavage predictions, NNs outperformed all other classifiers (Figure 7). The optimized hyperparameters for NNs with one-hot encoding were Adam solver, rectifier (ReLU) activation, 1e-8 regularization, no oversampling, and 1 hidden layer with 10 neurons. Combining networks into ensembles again improved accuracy and stability, so the final results were generated with 5 replicates of 10-fold cross-validated (CV) networks with an average Matthews correlation coefficient (MCC) of 0.965. Very few false positives existed for any prediction method, but two putative sites were somewhat conserved within the Semliki Forest (SF) complex nsP1 MTase-GTase core.[55] These two sites are predicted to be ordered and not solvent exposed and so are likely not biologically important. When applied to the human proteome, 714 of 20,350 proteins were predicted to be cleaved at least once. Enrichment analysis did not return useful results as for coronavirus protease predictions, so this large list was instead reduced as described in Methods to discuss only the most likely meaningful cleavages.

**Figure 7:**
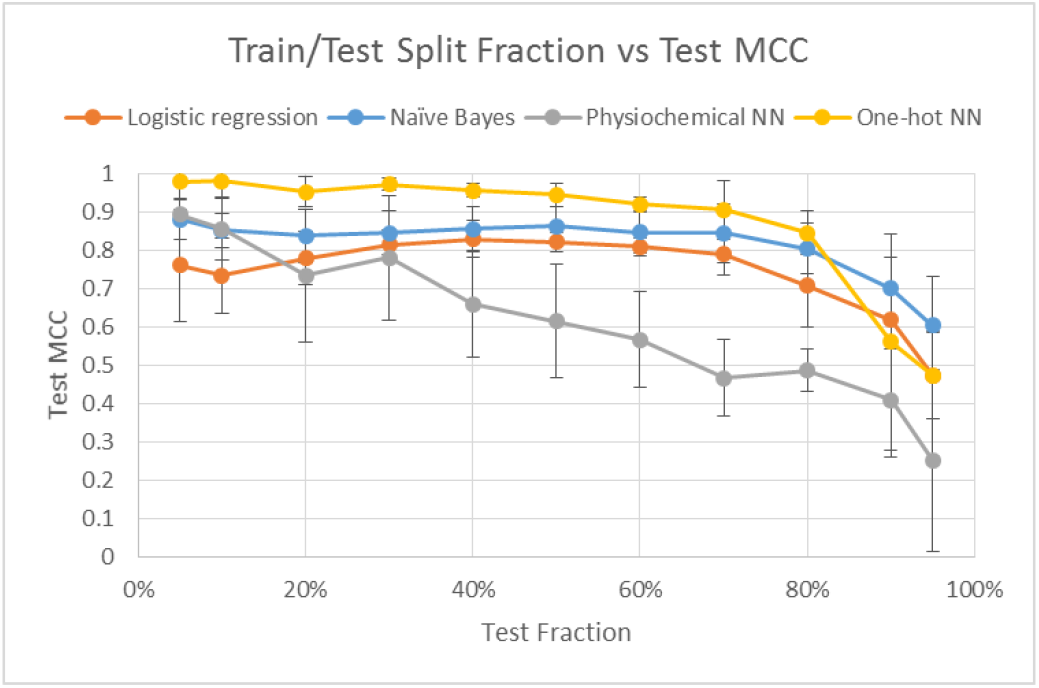
Train/test split fraction versus MCC demonstrating that the entire data set is not required for satisfactory accuracy.

## Discussion

Similar biases and caveats exist for nsP2pro predictions as with previous viral protease predictions,[14, 15] yet many host pathways likely perturbed by nsP2pro are discoverable. Experimental validation is, however, required for all of the following hypotheses. Reviewing only predicted cleavages with the highest scores and sufficient SASAs and relevant tissue expression produced a more targeted list of cleavages for interpretation. This list contains many proteins involved in the cytoskeleton and extracellular matrix (ECM), protease inhibition, protein translation/folding/glycosylation/ubiquitination, cellular differentiation including the transforming growth factor beta (TGF-β) and tumor necrosis factor alpha (TNF-α) pathways, inflammation, and vesicle trafficking.

Cleavage of many cytoskeletal proteins likely contributes to altered virus and host component trafficking, yet cytoskeletal drugs have remained mostly ineffective against alphaviruses. These drugs may help prevent initial endocytosis,[56, 57] internalization of replication complexes in spherules into cytopathic vacuoles,[58] and virion release, yet alphaviruses are known to still replicate in their presence.[59, 60]

In the extracellular matrix, the most obviously affected region in arthritis, nsP2pro likely cleaves many structural proteins. In particular, cleavage of lubricin’s hemopexin-like domain, similar to its normal cleavage by a human subtilisin-like proprotein convertase (SPC), may disrupt its ability to bind other proteins at the cartilage surface, reducing lubrication and promoting inflammation.[61] Cleavage of collagen alpha-1(XII) may disrupt normal shock-absorbing function.[62] In addition to the general matrix disruption likely resulting from cleavage of perlecan, elastin, von Willebrand factor A domain-containing protein 1 (WARP),[63] laminin, nidogen-1,[64] and thrombospondin-3 and −4,[65] degradation products of elastin and perlecan (laminin-like globular domain (LG3) of endorepellin) are documented to promote joint inflammation[66] and prevent angiogenesis in avascular cartilage,[67] respectively. Cleavage of specifically laminin subunit gamma-1 would not disrupt laminin binding membrane-bound integrins and dystrophins or extracellular collagen, but it may disrupt gamma subunit binding to nidogen and polymerization,[68, 69] freeing it up to be a more accessible alphavirus receptor.[70, 71] Similarly extracellularly secreted although not structural, nsP2pro has predicted cleavages near and within the bait regions of the serum and synovial fluid antiproteases alpha-2-macroglubulin (A2M) and pregnancy zone protein (PZP).[72, 73]

Noteworthy cleaved proteins involved in translation include signal recognition particle receptor subunit alpha (SRPRA), eukaryotic peptide chain release factor GTP-binding subunits (ERF3A/B), and La-related protein 1 (LARP1). Cleavage of SRPRA between its N-terminal SRX domain and its C-terminal targeting complex may reduce translocation of many proteins into the endoplasmic reticulum (ER). Only the viral structural polyprotein contains a signal peptide, so this cleavage may also relate the kinetics of its N-terminal capsid protein autocleavage to its subsequent SRP association.[74, 75] Cleavage of release factors ERF3A/B expressed in osteoblasts and osteoclasts may promote stop codon readthrough critical to alphavirus infection.[76, 77] Cleavage of LARP1 may have effects similar to its documented inhibition by alphavirus capsid protein or by mTOR often activated early in infection.[78] This mechanism of inhibiting host translation differs between mosquito and vertebrate cells, is not required for viral production, but may be required for internalization of the replication complex.[79] Unlike viruses requiring eukaryotic initiation factor 2 (eIF2) for translation, alphavirus downstream hairpin loop (DLP)-mediated translation does not benefit from amino acid starvation, so the effects of alphaviruses on mTOR are more straightforward than those of picornaviruses, flaviviruses, etc. For example, cleavage of the E3 ubiquitin ligase tetratricopeptide repeat protein 3 (TTC3) may help activate AKT and downstream protein synthesis similar to its direct interaction with nsP3,[79, 80] and cleavage of TRIM63 may stabilize many proteins against amino acid starvation-associated degradation.

Also after translation, cleavage of the immunophilins peptidyl-prolyl cis-trans isomerases (PPI) H and FKBP10 may affect folding of many relevant proteins. These may (1) have immunosuppressive effects similar to inhibition by tacrolimus, (2) modulate calcineurin, ribonuclease A, and some interleukins with cis-prolines in their native states, (3) affect proline- and hydroxyproline-rich collagen structure (supported by nsP2pro’s cleavage of prolyl 4-hydroxylase subunit alpha-1 and by documented excretion of proline and hydroxyproline in the urine of CHIKV-infected patients),[81] and (4) modulate alphavirus nsP3 proline-rich domain binding to amphiphysins involved in membrane bending of alphavirus-induced membrane organelles.[82, 83]

Enzymes involved in glycosylation also have predicted cleavages, although the differential effects this would have on viral versus host protein glycosylation remain unknown. Cleavage of mannosyl-oligosaccharide glucosidase (MOGS) may broadly disrupt viral protein glycosylation as in congenital disorders of this enzyme,[84] and cleavage of beta-1,4-galactosyltransferase 3 (B4GALT3) may disrupt complex N-linked glycans on immunoglobulins as in RA.[85] Cleavage of ER degradation-enhancing alpha-mannosidase-like protein 2 (EDEM2)[86] may disrupt host ERAD of viral glycoproteins[87] or redirect viral glycoproteins away from the cell membrane for internal budding as in SINV-infected mosquito cells.[88, 89] Cleavage of phosphoacetylglucosamine mutase (PAGM) may contribute to disruption of glycosaminoglycan polymers in cartilage and contribute to arthritic symptoms,[90] however N-acetylglucosamine (GlcNAc) supplementation sometimes used to treat osteoarthritis (OA) may be counterproductive given (1) it makes up some alphavirus glycans,[91] (2) it promotes replication of many other viruses *in vitro* and *in vivo*,[92] and (3) O-linked GlcNAc glycosylation of p65 aggravates TNF-α-mediated inflammation in rheumatoid arthritis (RA).[93]

Other predicted cleavages involved in protein degradation include the SUMO-specific E1 enzyme SAE1, the E2 enzyme UBE2Q1, the E3 enzyme UBR4, ubiquitin-1, and tripeptidyl peptidase 2 (TPP2). As with other viruses, modulation of SUMOylation is nontrivial and likely time-dependent; depletion of the only E2 for SUMO, UBC9, protects against CHIKV infection in mice,[94] yet depletion of SUMOylation enhances SFV replication in mosquito cells.[95] Cleavage of UBE2Q1 in muscle may disrupt B4GALT1-mediated cell adhesion to laminin and promote myoblast and satellite cell differentiation and syncytia formation[96, 97] to allow infection of myofibers without virion egress.[98–100] This is supported by the ability of alphaviruses to form filopodia-like protrusions mediating cell-to-cell transmission[101] and possibly to shield the virus from antibodies, making effective vaccination more difficult.[102] Cleavage of UBR4 may disrupt the N-end rule[103] to stabilize an inhibitor of apoptosis as with a picorna-like virus[104] or to stabilize cleaved viral functional proteins with less stable N-termini.[105] Cleavage of ubiquilin-1 between its ubiquitin-associated (UBA) and ubiquitin-like (UBL) domains may disrupt its trafficking ubiquitinated proteins to the proteasome[106] or its targeting of transmembrane proteins.[107, 108] Cleavage of the proteolytic TPP2 downstream of the 26S proteasome may promote viral susceptibility as in TRIANGLE disease[109] and may affect the pool of short peptides available for MHC class I presentation.[110]

TGF-β is known to be elevated in RA and during alphavirus infection,[111] and its inhibition can reduce joint swelling yet does not reduce viral titer[112] and can even promote CHIKV-mediated cell death *in vitro*.[113] Cleavage of latent TGF-β binding protein 3 (LTBP3) near its ECM-binding C-terminus may be one mechanism alphaviruses employ to encourage release of TGF-β from latency-associated peptide (LAP) when combined with normal cleavage by host proteases in its N-terminal hinge and C-terminus regions.[114, 115] These pathways are counterintuitive due to the many differential effects of TGF-β and bone morphogenetic proteins (BMPs) on different cell types and their interactions throughout their differentiation.[116] In the mesenchymal (MSC) lineage, TGF-β stimulates proliferation and differentiation of MSCs into chondrocytes and osteoblast progenitors into osteoblasts with downregulated RANKL. The closely related BMPs, however, can oppose TGF-β and are required for late stage osteoblast differentiation through their different SMAD signal transducers.[117] In the hematopoietic (HSC) lineage, TGF-β keeps HSCs in hibernation and prevents osteoclast progenitor differentiation into mature osteoclasts at least partially due to disrupted RANKL/OPG ratio.[118] Alphavirus infection is, however, more complex than elevated TGF-β alone and is associated with increased RANKL/OPG ratio and therefore osteoclastogenesis[119, 120] likely via upregulated IL-6 positive feedback[121, 122] and disrupted osteoblastogenesis via reduced RUNX2.[123] This TGF-β disruption may direct stem cells toward differentiated lineages more susceptible to infection, yet disruption of other pathways may be able to prevent apoptosis in these differentiated cells. For example, cleavage of TNFR2 may prevent TNF-α transduction and even shed soluble TNFR2 which can, like its alternately spliced products, antagonize its full-length activity and downstream apoptosis of infected cells.[124]

Likely also in an attempt to prevent inflammation and death of infected cells, cleavage of PYCARD between its pyrin domain (PYD) and caspase recruitment domain (CARD) may disrupt normal inflammasome formation and act like host CARD only proteins (COPs) or pyrin only proteins (POPs), inhibiting caspase 1 and its target cytokines.[125] As for other inflammatory molecular classes likely affected, polyamines are generally required by RNA viruses,[126, 127] and exogenous polyamines can restore inflammation and immune dysregulation in RA and OA.[128–130] Cleavage of diamine acetyltransferase 1 (SAT1) would increase intracellular polyamine concentration by preventing export and may (1) promote DNA methylation and reduce host transcription via S-adenosyl methionine (SAM) metabolism,[131] (2) downregulate IL-2 in PBMCs and contribute to decreased T cell effector function as in RA,[132] (3) enhance translation of polyproline motifs such as collagen and alphavirus nsP3 proline-rich domain,[133] (4) speed up peptidyl-tRNA hydrolysis by termination factor eRF1 via its hypusine modification,[134] and (5) promote stop codon readthrough by altering tRNA conformation.[135] Cleavage of histidine decarboxylase (HDC), glutathione hydrolase 5 (GGT5), and phospholipase A and acyltransferase 3 (PLAAT3) would, however, oppose typical inflammation in RA (where histamine[136] and lysophospholipids[137] are elevated and glutathione[138] is depleted), serving as a reminder that viral arthritis is caused more by the immune response to infection than by the virus or nsP2pro.

Lastly, the effects of predicted cleavages in vesicle transporting proteins are difficult to interpret because both retrograde and anterograde pathways are affected. Major differences exist between alphavirus-induced mammalian and mosquito membrane rearrangements, so experimental characterization is required to understand the relevance of these cleavages in each host.[139, 140]

## Conclusion

These predicted cleavages hint at many expected and novel mechanisms and indicate that nsP2pro is a much more important virulence factor than previously believed. Substrate docking and molecular dynamics may provide additional information about molecular mechanisms of these proteases, and protease-specific kinetics and biological significance of these cleavages require experimental verification. Expansion of this data set to include all of *Martellivirales* or *Alsuviricetes* may also provide insight into how these molecular mechanisms evolved, but their inclusion into a cleavage prediction training data set would likely worsen the trained model’s accuracy for the human viruses discussed here. Even though many caveats exist without experimentation, similar prediction and interpretation should be performed for all other viral proteases.

